# Homocysteine-induced endoplasmic reticulum stress activates FGF21 via CREBH, resulting in browning and atrophy of white adipose tissue in *Bhmt* knockout mice

**DOI:** 10.1101/2021.09.08.459498

**Authors:** Manya Warrier, Evan M. Paules, Walter B. Friday, Frances Bramlett, Hyunbae Kim, Kezhong Zhang, Isis Trujillo-Gonzalez

**Author notes:** Correspondence:* Isis Trujillo-Gonzalez, PhD, Research Assistant Professor, UNC Nutrition Research Institute, University of North Carolina at Chapel Hill, 500 Laureate Way, Room 2018, Kannapolis, NC 28081.

## Abstract

Betaine-homocysteine methyltransferase (BHMT) catalyzes the transfer of methyl-groups from betaine to homocysteine (Hcy) producing methionine and dimethylgycine. In this work, we characterize *Bhmt* wildtype (WT) and knockout (KO) mice that were fully backcrossed to a C57Bl6/J background. Consistent with our previous findings, *Bhmt* KO mice had decreased body weight, fat mass and adipose tissue weight compared to WT. Histological analyses and gene expression profiling indicate that adipose browning was activated in KO mice and contributed to the adipose atrophy observed. BHMT is not expressed in adipose tissue but is abundant in liver, thus, a signal must be originating from the liver that modulates adipose tissue. We found that, in Bhmt KO mice, homocysteine-induced endoplasmic reticulum (ER) stress, with activation of hepatic transcription factor cyclin AMP response element binding protein (CREBH), mediated an increase in hepatic and plasma concentrations of fibroblast growth factor 21 (FGF21), which is known to induce adipose browning. CREBH binds to the promoter regions of FGF21 to activate its expression. Taken together, our data indicate that deletion of a single gene in one-carbon metabolism modifies adipose biology and energy metabolism. It would be interesting to determine whether people with functional polymorphisms in *BHMT* exhibit a similar adipose atrophy phenotype.

## INTRODUCTION

Betaine-homocysteine *S*-methyltransferase (BHMT) is an important Zn-dependent thiol-methyltransferase that catalyzes the formation of methionine from homocysteine using betaine as its methyl donor (1, 2). Methionine is subsequently converted to *S*-adenosylmethionine (SAM) and is used for various methylation reactions (3). BHMT is one of the most abundant proteins in the liver, amounting to 0.6-1% of total protein (4), and it is also found in kidney, the eye lens, and at lower activities in other tissues, but not in adipose (5, 6). Mice in which *Bhmt* was deleted (whole body; *Bhmt* KO) have increased hepatic concentrations of the substrates betaine and homocysteine (Hcy) (5, 7). These KO mice develop increased energy expenditure associated with lower body weight compared to their wild type (WT) littermates and develop lipodystrophy and fatty liver (5, 7). At 1 year of age, 64% of *Bhmt* KO mice develop hepatic tumors (5, 7). Though the mechanisms underlying the hepatocarcinogenesis have been explored (3), those involved in the adipose wasting have not been addressed, we do so in this paper.

In this study, we show that deletion of *Bhmt* in mice causes increased Hcy concentrations in tissues, and that this initiates a signaling cascade involving endoplasmic reticulum stress (ER stress) with activation (cleavage) of cyclic AMP response element binding protein H (CREBH) generating a transcription factor that promotes the expression of genes including fibroblast growth factor 21 (*Fgf21*; previously, we reported increased Fgf21 concentrations produced by the liver in *Bhmt* KO mice (7)). Fgf21 stimulates adipose browning and energy expenditure by upregulating the expression of the transcriptional co-activator peroxisome proliferator-activated receptor gamma coactivator **1**-alpha (Pgc-1α), as well as uncoupling protein 1 (Ucp1). This culminates in adipose wasting in *Bhmt* KO mice.

## Results

### Deletion of Bhmt promotes adipose atrophy in fully backcrossed mice

We previously reported that *Bhmt* knockout mice on a mixed 129/SV x C57BL/6J background (generations F3-F5), between 7-12 weeks of age, had reduced adipose mass and smaller-sized adipocytes (7). Since genetic background of mice can have a profound influence on the metabolic phenotype of mice (8), we decided to reexamine this lipodystrophy phenotype after backcrossing *Bhmt* KO mice to C57Bl/6 to generate a near congenic (99.74%) line. In this near congenic line, we confirmed that, in the *Bhmt* KO compared to wild type (WT), there was a significant reduction in total body weight (**Fig 1A**) and adipose weight **(Fig 1B)** in mice. Histological analysis of adipose tissue taken from *Bhmt* KO mice showed reduced adipocyte cell size and reduced size of lipid droplets in both gonadal white adipose tissue (gWAT; data not shown) and inguinal white adipose tissue (iWAT) as compared to WT **(Fig 1C and Fig 1D)**. Adipose atrophy is characterized by reduced fat/lean mass and the ‘slimming of adipocytes’ in both size and volume (9), and our data show that this process was dependent on *Bhmt* status.

**FIGURE 1.**
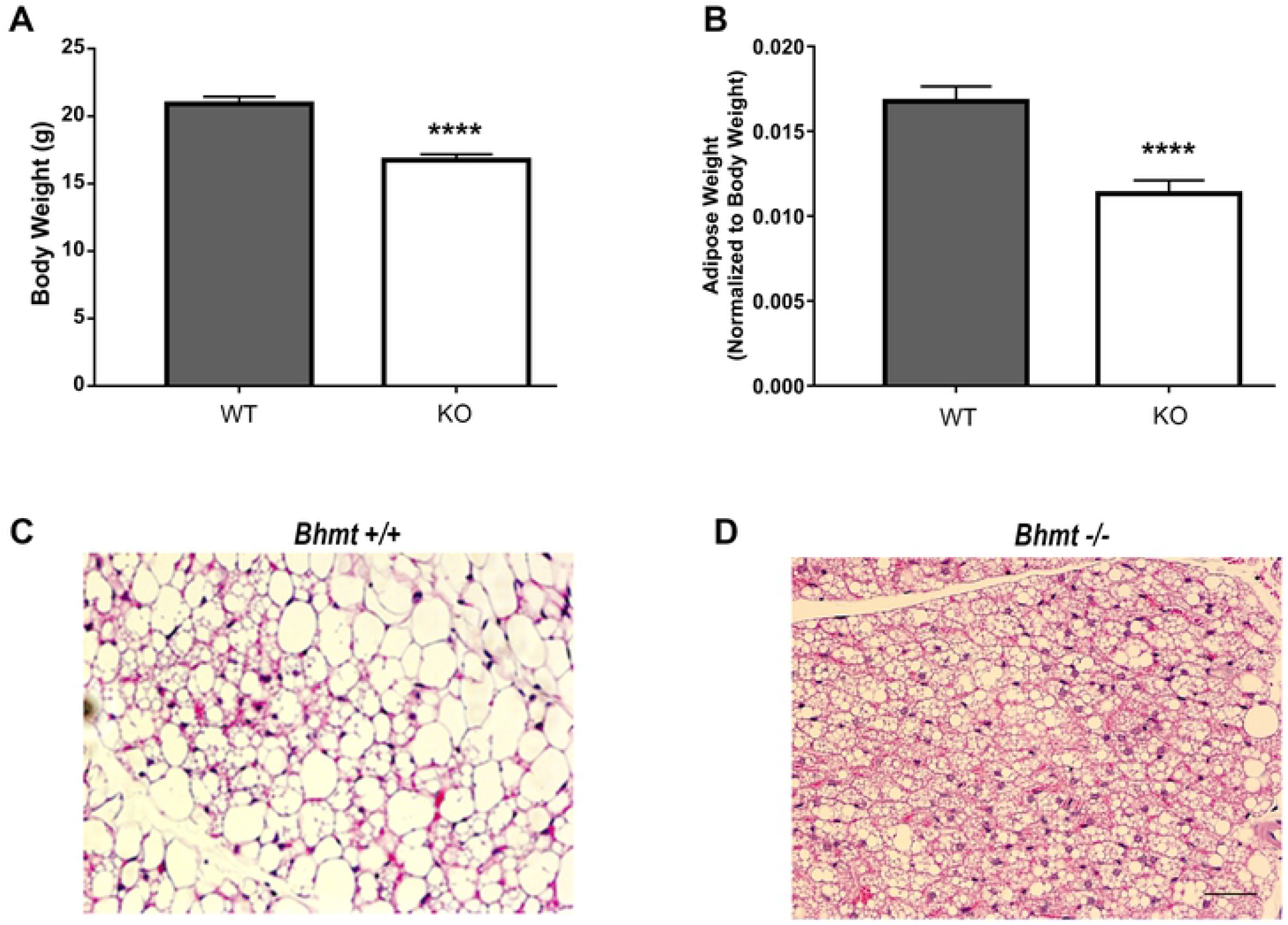
Lack of the *Bhmt* gene induces adipose atrophy in mice. **(A)** Bodyweight loss of Bhmt knockout (*Bhmt-KO*) compared to Bhmt wild type (*Bhmt-WT)*. **(B)** Adipose weight normalized over body weight. *n*= 25 *Bhmt-KO; n*= 24 *Bhmt-WT*. **(C) and (D)** Representative stainings of sections from inguinal white adipose tissue (iWAT) from *Bhmt-WT* **(C)** and *Bhmt-KO* **(D)** Scale bar= 50 mm. Results represent mean ± SEM. ****P≤0.0001 by unpaired t-test.

### *Adipose atrophy in Bhmt* KO *mice is associated with adipose browning in inguinal adipose depots*

Since smaller adipocytes and increased whole body energy expenditure and heat production are classical features of browning of WAT (10), and since adipose browning is known to promote adipose atrophy in several mouse models (10-14), we sought to determine whether the WAT atrophy observed in *Bhmt* KO mice is due to WAT browning. We first measured a number of molecular markers that are frequently associated with adipose browning (15-17). Uncoupling Protein 1 (*Ucp1)* mRNA **(Fig 2A)** along with mRNA for other thermogenic genes such as Peroxisome proliferator-activated receptor gamma coactivator 1-alpha (*Pgc-1α)* and the lipid-droplet-associated protein cell death-inducing DFFA-like effector A (*CideA*) **(Fig 2B – C)** were all significantly upregulated in iWAT collected from *Bhmt* KO mice compared to WT. Together, these results indicate that lack of *Bhmt* is sufficient to induce the expression of adipose browning markers.

**FIGURE 2.**
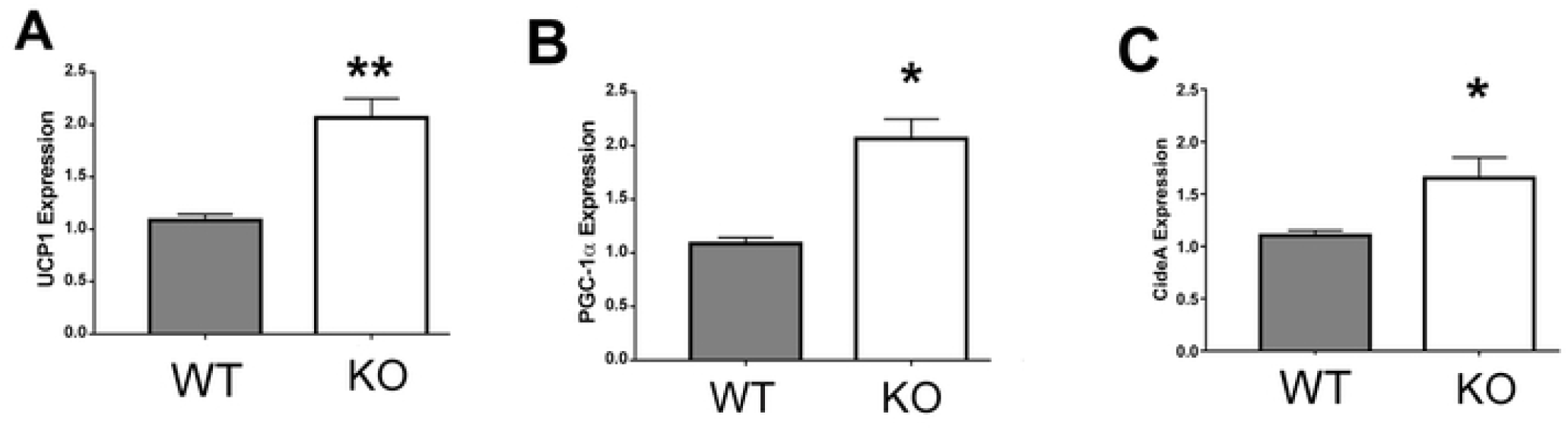
Lack of the *Bhmt* gene induces the expression of beige remodeling markers that induce browning. mRNA levels of beige remodeling markers *Ucp1* **(A)**, *Pgc1a* **(B)**, and *Cidea* **(C)** in inguinal adipose tissue (iWAT) of *Bhmt-WT* and *Bhmt-KO* mice. Relative quantitative values (normalized to 36B4) are reported as fold change. Results represent mean ± SEM. *P≤0.05, **P≤0.01 by Mann-Whitney test (A and C) and by unpaired t-test (B and D). *n* = 10 per group.

### *Bhmt* KO *livers have increased homocysteine concentrations*

Plasma total Hcy concentrations were significantly increased in *Bhmt* KO mice on a mixed 129/SV x C57BL/6J background (generations F3-F5), as we previously reported (7). We now show that, in the fully backcrossed *Bhmt* KO mice, both plasma Hcy concentrations (∼11 fold) and liver Hcy concentrations (∼2 fold) were increased in KO as compared to WT mice (**Fig 3A and B**). Thus, loss of *Bhmt* results in the accumulation of plasma and liver Hcy.

**FIGURE 3.**
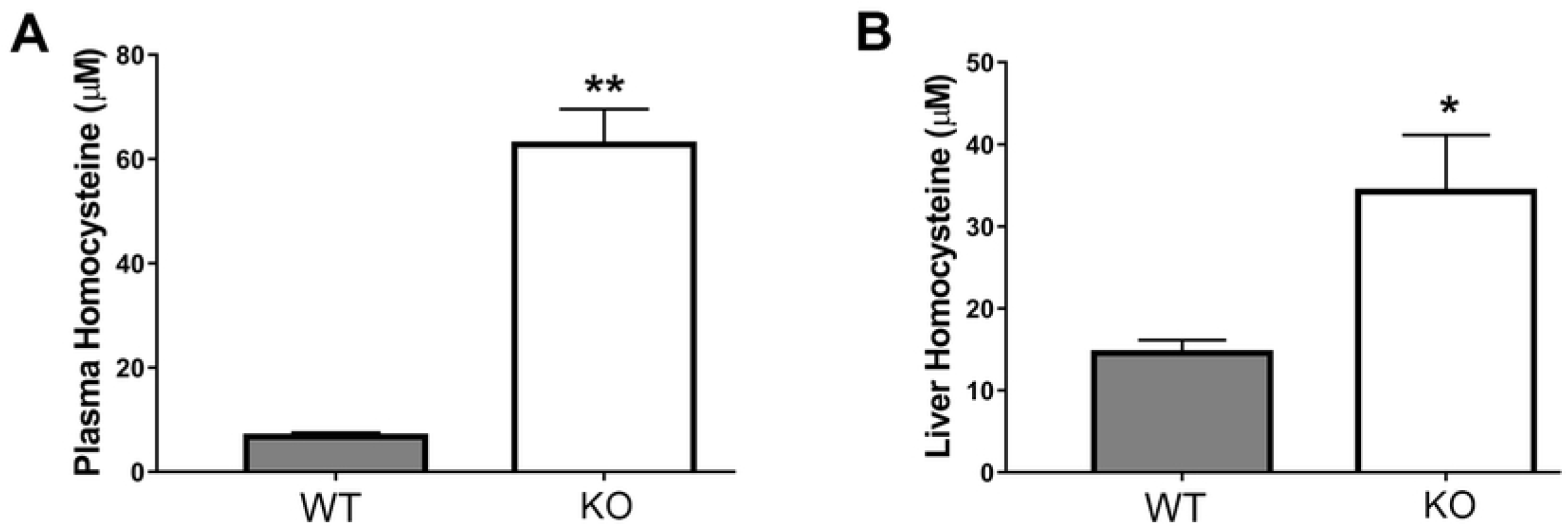
Increase in plasma and liver homocysteine (Hcy) levels in *Bhmt-KO* mice. **(A)** Plasma levels of Hcy are increased in *Bhmt-KO* mice ∼50 fold when compared with *Bhmt-WT* mice. *n*=5 per group. **(B)** Liver Hcy levels were also increased ∼20 fold in *Bhmt-KO* mice when compared with *Bhmt-WT. n*= 7 *Bhmt-KO; n*=5 *Bhmt-WT*. Results represent mean ± SEM. *P≤0.05, **P≤0.01 by Mann-Whitney test.

### *Bhmt* KO *livers have increased ER stress and have more activated CREBH*

Since high tissue Hcy is a known cause of ER stress, which in turn regulates a number of transcription factors residing in the ER (18, 19), we decided to see if *Bhmt* KO mice experience increased ER stress compared to WT. We measured gene expression of DNA damage-inducible transcript 3, also known as C/EBP homologous protein (CHOP) and Activating Transcription Factor 3 (ATF3), as indicators of ER stress (20-30). We found that expression of these genes were increased by 1.5-fold and 3-fold, respectively, in *Bhmt* KO compared to WT mouse livers (**Figs 4A and 4B**). Thus, *Bhmt* KO mice have increased ER stress compared to their WT counterparts.

**FIGURE 4.**
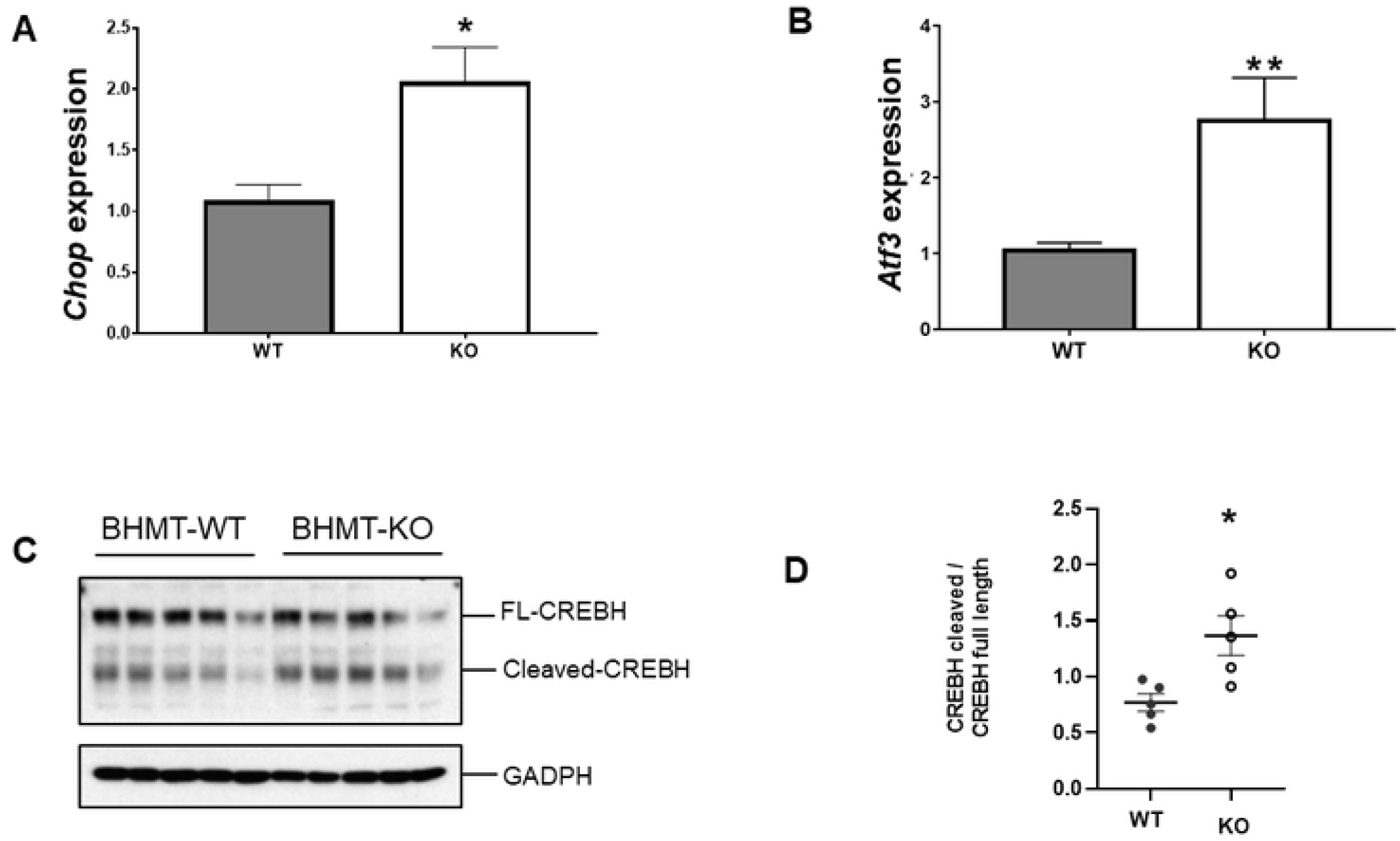
Endoplasmic reticulum (ER) stress is increased in *Bhmt-KO* livers and exhibit activation of CREBH. mRNA levels of ER stress markers *Chop* **(A)** *n*= 10 per group, and *Atf3* **(B)** are increased in liver *Bhmt-KO* mice compared to WT. *n*= 7 *Bhmt-KO; n*=8 *Bhmt-WT* Relative quantitative values (normalized to 36B4) are reported as fold change. Results represent mean ± SEM. *P≤0.05, **P≤0.01 by unpaired t-test (A) and Mann Whitney test (B). **(C)** Representative western blot of full-length CREBH and cleaved CREBH from *Bhmt-WT* and *Bhmt-KO*. GAPDH was used as a loading control. **(D)** The ratio of cleavaged CREBH divided by full-length CREBH. *P≤0.05 by unpaired t-test.

Next, we searched for transcription factors that reside in the ER and are produced in response to ER stress and which are also known to regulate FGF21. We found that the hepatic transcription factor known as Cyclin AMP Responsive Element Binding Protein – H (CREBH) fulfilled the above criteria (19, 31, 32). We measured full length and activated CREBH in the liver lysates prepared from both *Bhmt* WT and KO mice by Western blot analysis and found that the cleaved activated form of CREBH was significantly increased in *Bhmt* KO compared to WT liver **(Figs 4C and 4D**).

### *Bhmt* KO *livers have increased FGF21 concentrations*

Since activated CREBH binds to the *Fgf21* promoter and activates its transcription (33), we suggest that this explains our earlier finding that *Bhmt* KO mice on a mixed 129/SV x C57BL/6J background (generations F3-F5) had increased FGF21 concentrations (7). We now show that in fully backcrossed *Bhmt* KO mice, compared to WT, plasma and hepatic FGF21 concentrations were increased more than 2-fold (**Figs 5A and 5B**).

**FIGURE 5.**
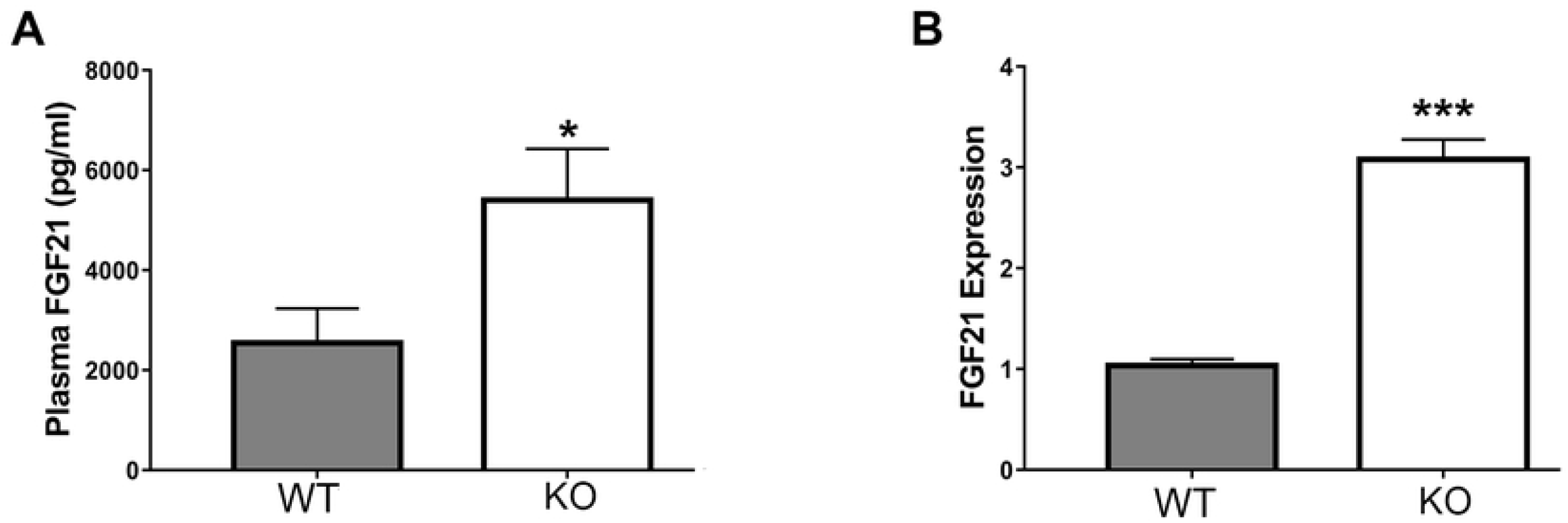
FGF21 is increased in *Bhmt-KO* plasma and liver. **(A)** Plasma FGF21 levels are increased in plasma from *Bhmt-KO* mice when compared to *Bhmt-WT. n=*9 per group Results represent mean ± SEM. *P≤0.05 by *t*-test. **(B)** mRNA levels of Fgf21 in the liver are increased ∼2 fold. Relative quantitative values (normalized to 36B4) are reported as fold change. ****P≤0.0001 by *t*-test. *n=*5 per group.

## Discussion

The deletion of *Bhmt* in mice results in the animal storing less fat in adipose tissue even though BHMT is not expressed in adipose tissue (7). This adipose atrophy is the result of reduced triglyceride storage within iWAT associated with increased energy expenditure and heat production as measured by indirect colorimetry without a matching increase in food consumption (7). We now report that the elevated Hcy concentrations that occur when the *Bhmt* gene is deleted, increase ER stress signalling, which results in generation of activated CREBH, and this, in turn, caused increased expression in hepatic FGF21. This FGF21 is transported via blood to adipocytes where it promotes the browning of white adipose tissue and increases expression of PGC-1α which increases mitochondrial number, and increases the expression of UCP-1 which uncouples mitochondrial respiration and thereby increases energy expenditure and heat production. (**Fig 6**).

**FIGURE 6.**
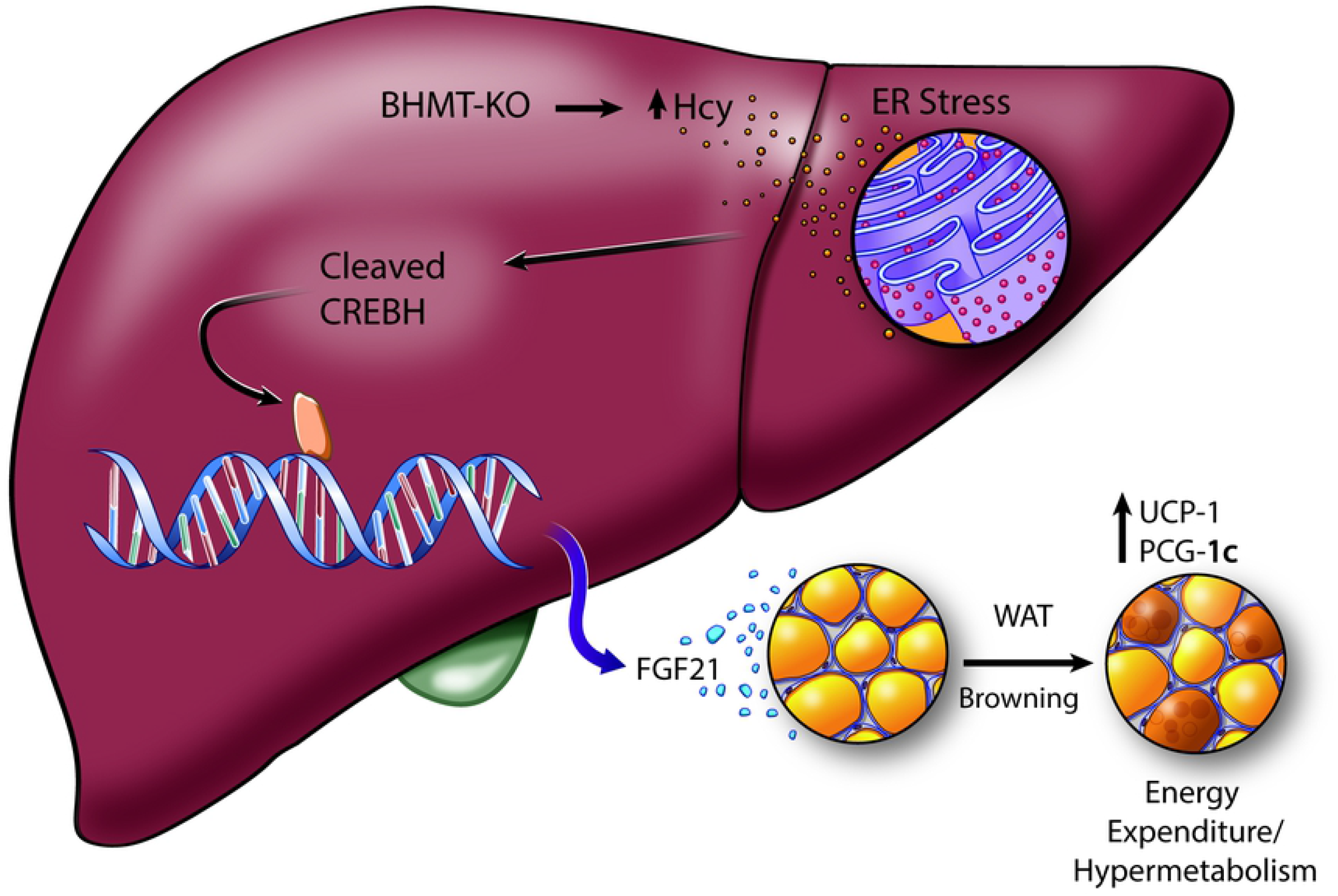
Schematic representation of the effects of the deletion of *Bhmt* in liver and iWAT. The schema summarizes our new findings where deletion of *Bhmt* in mice increases homocysteine levels leading to endoplasmic reticulum (ER) stress. ER stress led to an increase in the cleaved CREBH protein levels, which acts as a transcription factor that binds the FGF21 promoter. FGF21 high levels exert their effects in iWAT.

As noted earlier, BHMT catalyzes the formation of methionine from Hcy using betaine as its methyl donor (1, 2). As expected, deletion of *Bhmt* should increase concentrations of both substrates (betaine and Hcy) used by this enzyme. Increased concentrations of Hcy cause ER stress both *in vitro* and *in vivo* (20, 21, 23, 26, 27, 30, 34-36) by disrupting disulfide bond formation and thus leading to protein misfolding (21). Though we argue that it is the the accumulation of Hcy that results in ER stress and subsequent browning of adipose tissue, it is possible that the accumulation of the other precursor, betaine, also contributes to adipocyte browning as feeding mice a diet containing 5% betaine increases plasma concentrations of FGF21 (7, 37). Since *Bhmt* deletion resulted in reduced methylation potential by increasing *S*-adenosylhomocysteine concentrations, in earlier studies we examined whether the FGF21 promoter region might be hypomethylated in the *Bhmt* KO mouse, leading to increased expression of this gene. However, reduced representation bisulfite sequencing performed on liver DNA from WT and KO mice did not reveal any methylation differences in this gene (3).

ER stress is initiated by numerous metabolic stressors including high concentrations of homocysteine (20, 21, 23, 26) and has been associated with hepatic lipid accumulation, obesity and cancer (24, 38). Also, it has been implicated in WAT browning (39). Transcription factors that are regulated by ER stress include Sterol Regulatory Element Binding Proteins (SREBP) and CREBH (29). CREBH is ER-tethered and is synthesized in the liver as a precursor which then gets activated via cleavage by Golgi-localized proteases and the activated form then accumulates in the nucleus to act as a transcription factor (19) that promotes expression of the liver-secreted peptide endocrine hormone FGF21 (31, 33). Promoter analysis studies show that CREBH can bind and activate FGF21 promoter at position -60 to -40 bp. Chromatin immunoprecipitation studies reveal that CREBH directly binds to the FGF21 promoter and controls the expression and plasma levels of FGF21 (33, 40). Using CREBH KO and CREBH over-expression mouse models it has been shown that FGF21 mediates many of CREBH’s effects on fatty acid metabolism and ketogenesis (41). Also, FGF21 is responsible for the body weight loss induced by CREBH over-expression and particularly the fat mass reduction (31). Therefore, it is reasonable to propose that CREBH is the upstream transcriptional regulator of FGF21 in *Bhmt* KO mice. In addition to deletion of *Bhmt*, essential amino acid restriction, fasting and impaired muscular and hepatic autophagy induce ER stress and result in substantial increases in circulating FGF21 and UCP1 levels in adipose (42, 43).

Many endocrine and autocrine signals stimulate adipose browning, including FGF21 (44). FGF21 binds to its receptor (FGFR) and coreceptor β-Klotho (KLB) to activate a downstream signaling cascade that ultimately leads to expression of its target genes (43). FGF21 stimulates adipose browning and energy expenditure by upregulating the expression of transcriptional co-activator PGC-1α in adipose tissue (17, 43, 45). Browning of white adipose tissue is characterized by the appearance of brown-like or beige adipocytes within WAT (46, 47). These inducible beige adipocytes are morphologically similar to brown adipocytes and express uncoupling protein 1 and contribute to thermogenesis (39, 47).

As noted earlier, adipose atrophy is characterized by reduced fat/lean mass and the excessive ‘slimming of adipocytes’ in both size and volume (9). Increased metabolic rate and adipose browning has been proposed as causes for adipose atrophy (9, 11-14, 48-52). Even though browning of WAT is generally considered beneficial in obesity (reducing body weight and increasing energy expenditure), several lines of evidence suggest that it also is associated with adverse outcomes such as hepatic steatosis, cancer associated cachexia (CAC) and burn-related cachexia (11, 14, 49, 50).

Is reduced BHMT expression likely to be a problem in people? Several functional *Bhmt* variants have been identified in humans which are associated with increased risk for cancer and other diseases (53-56), however no information is available on the metabolic phenotype of humans carrying those variants. It would be interesting to explore whether people with functional *Bhmt* variants have a metabolic phenotype similar to that which we describe in mice, and determine whether proposed Hcy-CREBH-FGF21–adipose browning pathway drives this phenotype. This would not only help us to understand how genetic variations in one carbon metabolism affect obesity but also our understanding of how adipose atrophy develops in diseases such as cancer.

## MATERIALS AND METHODS

### Animals

Mice used in these experiments were bred and maintained at the David H. Murdock Research Institute (DHMRI), Center for Laboratory Animal Science facilities. All animal experiments were performed in accordance with the protocols approved by David H. Murdock Research Institute Institutional Animal Care and Use Committee. The study was carried out in compliance with the ARRIVE guidelines.

*Bhmt* KO mice were generated as previously described (5). *Bhmt* KO mice were fully backcrossed to C57B1/6 wild-type mice to generate a near congenic (99.73 %) mouse line. Genotyping of *Bhmt* animals was performed using the following primers: *Bhmt* WT_F 5’– GACTTT TAAAGAGTGGTGGTACATACCTTG-3’, *Bhmt* WT_R -5’ – TCTCTCTGCAGCCACATCTGAACTTGTCTG-3’, *Bhmt* KO_F-5’ – TTAACTCAACATCACAACAACAGATTTCAG -3’, *Bhmt* KO_R 5’ –TTG TCGACGGATCCATAACTTCGTATAAT -3’. *Bhmt* WT and KO mice were mated and maintained *ad libitum* on a AIN 76A diet (Dyets, Bethelehem, PA, USA) and were kept in a temperature-controlled environment at 24°C and exposed to a 12 hours light and dark cycle. At 6-8 weeks, mice were euthanized and tissue collection was performed.

### Histological analysis

Tissues were fixed in buffered formalin, dehydrated in ethanol and then transferred to xylene solution for embedding in paraffin. Serial sections at 5 mm thickness were made from paraffin-embedded tissue and then stained with hematoxylin and eosin. Images were analyzed with light microscopy. Adipocyte area was calculated by measuring the area of cells per condition, at 200x magnification, using Image J, and presented as mean ± SEM.

### RT-PCR analysis

Total RNA was extracted from tissues of *Bhmt* WT and *Bhmt* KO mice, using RNAeasy mini Kit (Qiagen, Hilden, Germany). cDNA synthesis was performed by using a Script™cDNA SuperMix (Quanta BioSciences, Gaithrsburg, MD, USA). For quantitative real-time assays, amplification was performed by using PerfeCTa qPCR FastMix (Quanta Biosciences). We designed primers (Sigma) as follows: ***UCP1*** forward primer: ACTGCCACAACCTCCAGTCATT, reverse primer CTTTGCCTCACTCAGGATTGG; **PGC1a** forward primer AGCCGTGACCACTGACAACGAG, reverse primer GCTGCATGGTTCTGAGTGCTAGG; ***CIDEA*** forward primer: GCAACCAAAGAAATGCGGAATAG, reverse primer: CTCGTACATCGTGGCTTTGA; ***CHOP*** forward primer CAGCGACAGAGCCAGAAT; ***ATF3*** forward primer GAGGCGGCGAGAAAGAAA, reverse primer CACACTCTCCAGTTTCTC. Ct values were calculated by SDS 1.2 software (Applied Biosystems, Foster City, CA, USA) and normalized to *TATA* binding Ct values and expressed as 2 ^−(Ct(gene)-Ct (housekeeping gene))^.

### Western blot

Liver tissues were collected to evaluate CREBH levels. Protein extracts were preparared using RIPA lysis buffer (Sigma, ST. Louis, USA) supplemented with protease inhibitor cocktail (Complete, Roche) and sonicated. Total protein concentrations for all samples was quantified using BCA protein assay (Bio-Rad, Hercules, CA, USA). Proteins were loaded into SDS-PAGE gels and blotted on PVDF membranes. CREBH antibody was used at 1:1000 dilution. Enhancer chemiluminescence was used to detect protein. CREBH protein abundance was quantified using Image J (NIH, Bethesda, MD, USA). Data are presented mean ± SEM.

### FGF21 measurement

#### Serum

Blood samples from *Bhmt* WT and *Bhmt* KO mice, were collected and were subjected to centrifugation at 1000 g for 15 min at 4°C. **Liver**: Crushed liver samples were homogenized in cold phosphate-buffered saline (PBS) (Sigma) with protease inhibitors (Roche). Samples were subjected to centrifugation at 9,600 g for 15 minutes at 4° C. For both plasma and liver, supernantatant protein was quantified using BCA protein assay (Bio-Rad, Hercules, CA, USA) and diluted to equal concentrations before performing an enzyme-linked immunoabsorbent assay (ELISA) using a Mouse/Rat FGF21 Quantikine ELISA kit (R&D Systems, Minneapolis, MN)(57).

#### Homocysteine measurement

Plasma or liver was homogenized in dithiothreitol (DTT) and processed to dissociate the proteins by filtration, thereby extracting protein-bound Hcy. The protein-free filtrate was analyzed for total Hcy by liquid chromatography-electrospray ionization-tandem mass spectrometry (LC-ESI-MS/MS) as previously described (58, 59).

#### Statistical analysis

The number of samples per group are indicated in the figure legends. There were no experimental units or data points excluded. Statistical analyses were performed with Prism 7 (GraphPad Software, La Jolla, CA, USA). Data distribution was tested for statistical normality. The Brown-Forsythe test (F test) was used to compare group variances. Groups with equal distribution were compared using Students’ t test. Groups with unequal variances were compared using the nonparamentric Mann-Whitney test. Data are presented as means ± SEM.

## List of abbreviations

CIDEA: Cell death-inducing DFFA-like effector A
ATF3: Activating Transcription Factor 3
BHMT: Betaine Homocysteine-S-Methyltransferase
BAT: Brown adipose tissue
CHOP: c/EPB homologous protein
CREBH: Cyclic AMP response element binding protein
H FGF21: Fibroblast Growth Factor 21
Hcy: Homocysteine
PCG-α: Peroxisome Proliferator-activated receptor gamma coativator 1-alpha
SAM: S-adenosyl methionine
UCP1: Uncoupling Protein 1
WAT: White adipose tissue

## Acknowledgments

The authors thank Jennifer Owen (University of North Carolina at Chapel Hill, Nutrition Research Institute) for providing assistance with experiments; Dr. Steve Orena (University of North Carolina at Chapel Hill, Nutrition Research Institute) for providing metabolite services. This work was supported by U.S National Institutes of Health (NIH), National Institute of Diabetes and Digestive and Kidney Diseases Grants DK056350 and DK115380 (To Dr. Steven Zeisel).

## Author contributions

M. W. performed experiments, analyzed the data, conceived the study and wrote the manuscript, E. M. P. performed experiments, W. B. F. performed experiments, F.B. performed experiments, H. K. performed western blot experiments, K. Z. performed experiments, I. T-G. developed study designs, performed experiments, analyzed and interpreted the data, prepared figures and wrote the manuscript.

